## Supporting Information for "A phylogeny-informed characterisation of global tetrapod traits addresses data gaps and biases"

Mario R. Moura

Walter Jetz

**This PDF file includes:**

**Supplementary Figures**

**Supplementary Tables**

**Supplementary Figures**

Additional figures.

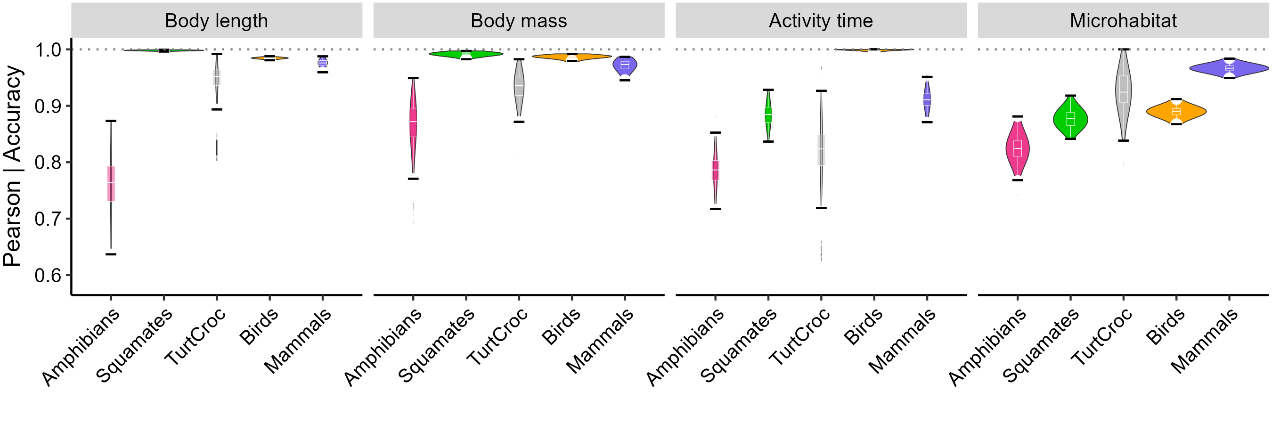

**Figure S1. Performance of phylogenetic multiple imputation across natural history traits of tetrapods.** The y-axis indicates the Pearson correlation coefficient for continuous traits (body length and body mass, log_10_ transformed) and the proportion of correctly classified entries (accuracy) for binary traits (types of activity time and microhabitat).

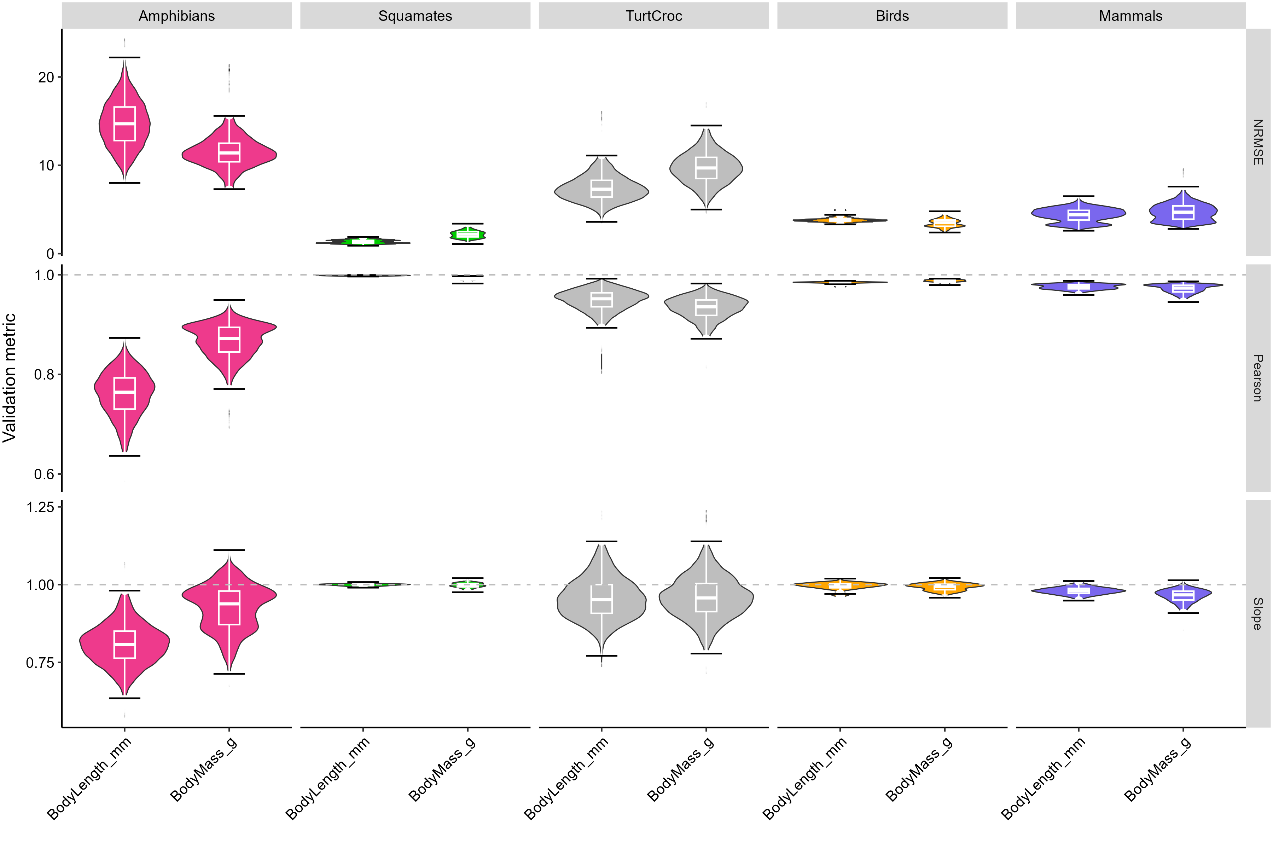

**Figure S2. Performance of phylogenetic multiple imputation for continuous attributes.** Validation metrics were computed between imputed and observed log_10_ attribute values, and include: NRMSE (Normalised Root Mean Square Error), Pearson (Pearson correlation coefficient), and Slope (linear regression slope).

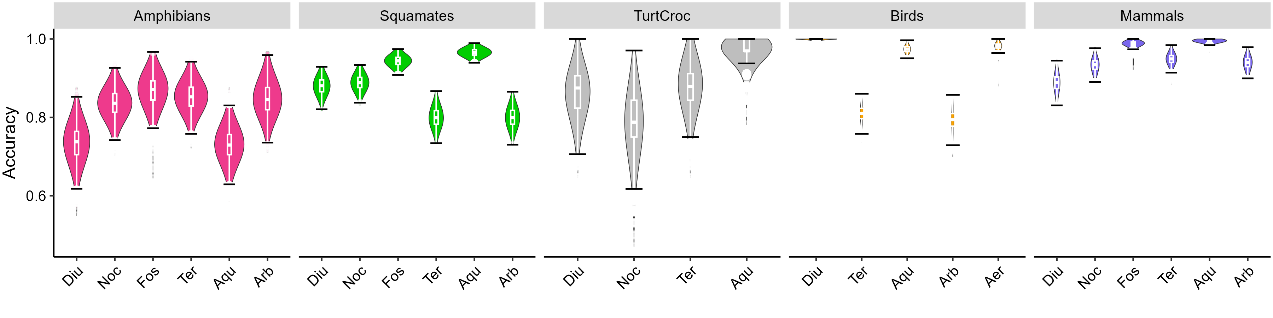

**Figure S3. Performance of phylogenetic multiple imputation in classifying activity time and microhabitat types.** The accuracy, measured as the proportion of correctly classified entries, was computed by comparing imputed and observed binary values. Results are reported separately for different types of activity time and microhabitat.

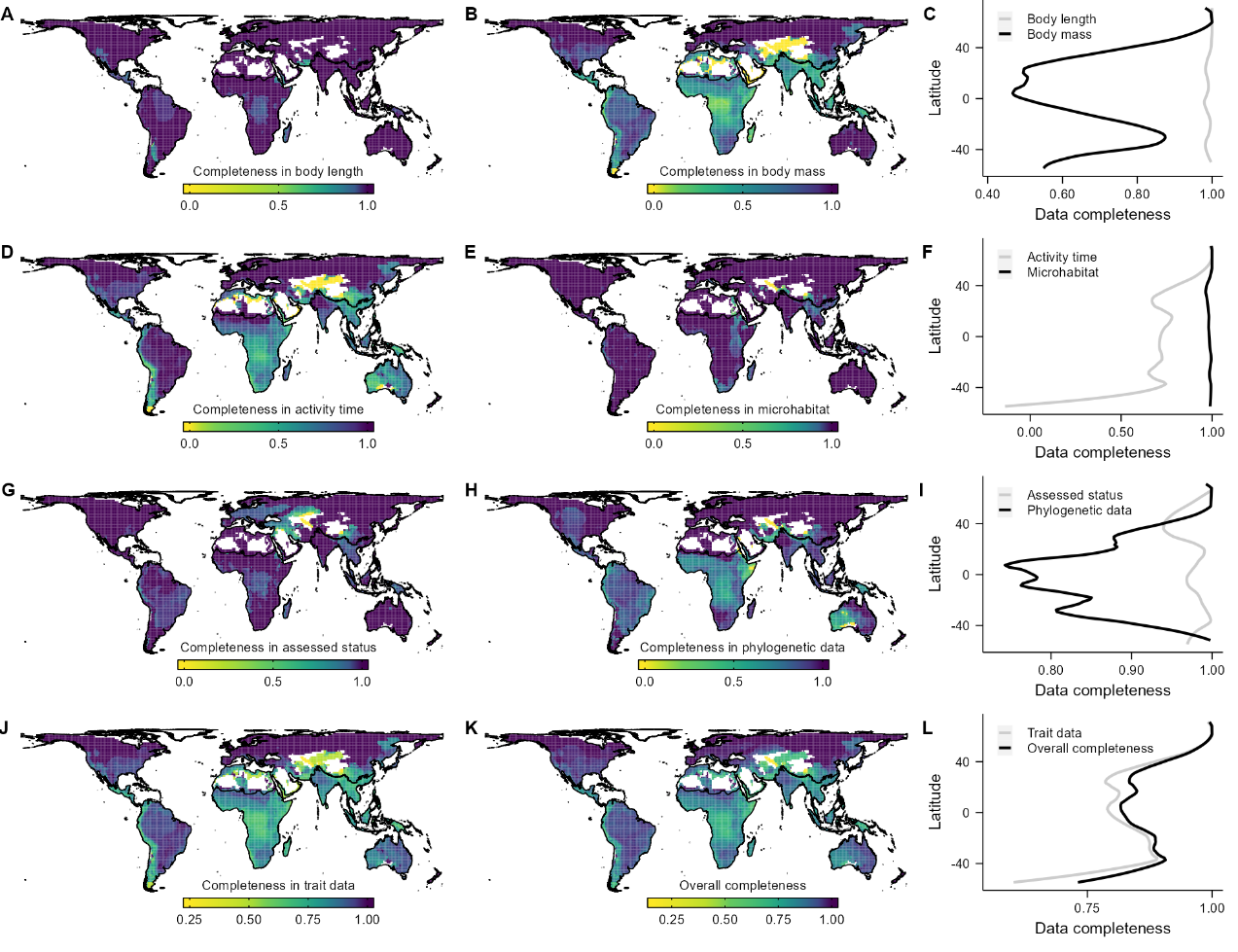

**Figure S4. Average data completeness across amphibian assemblages.** Proportion of species with observed values for: (A) body length, (B) body mass, (D) activity time, (E) microhabitat, (G) assessed threat status, (H) phylogenetic data, (J) attribute set, i.e., the average pattern for maps depicted in ABDE, and (K) the complete database, i.e., average pattern for maps depicted in ABDEGH. Maps show grid cell assemblages of 110 × 110 km size in an equal-area projection. Latitudinal plots (C, F, I, L) show the average values of these cells across latitudes. Colour ramps followed Jenks’ natural breaks classification.

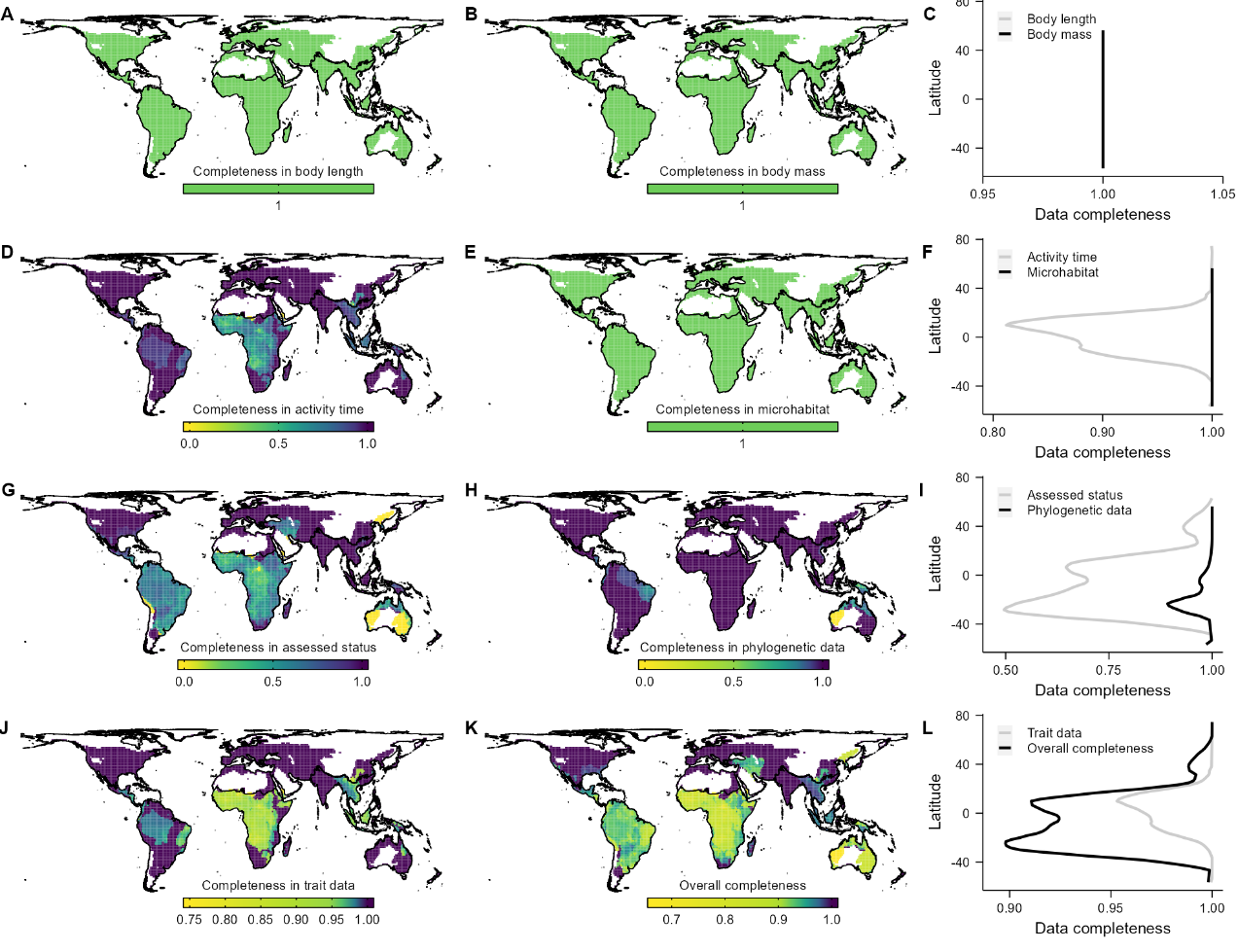

**Figure S5. Average data completeness across chelonian and crocodilian assemblages.** Proportion of species with observed values for: (A) body length, (B) body mass, (D) activity time, (E) microhabitat, (G) assessed threat status, (H) phylogenetic data, (J) attribute set, i.e., the average pattern for maps depicted in ABDE, and (K) the complete database, i.e., average pattern for maps depicted in ABDEGH. Maps show grid cell assemblages of 110 × 110 km size in an equal-area projection. Latitudinal plots (C, F, I, L) show the average values of these cells across latitudes. Colour ramps followed Jenks’ natural breaks classification.

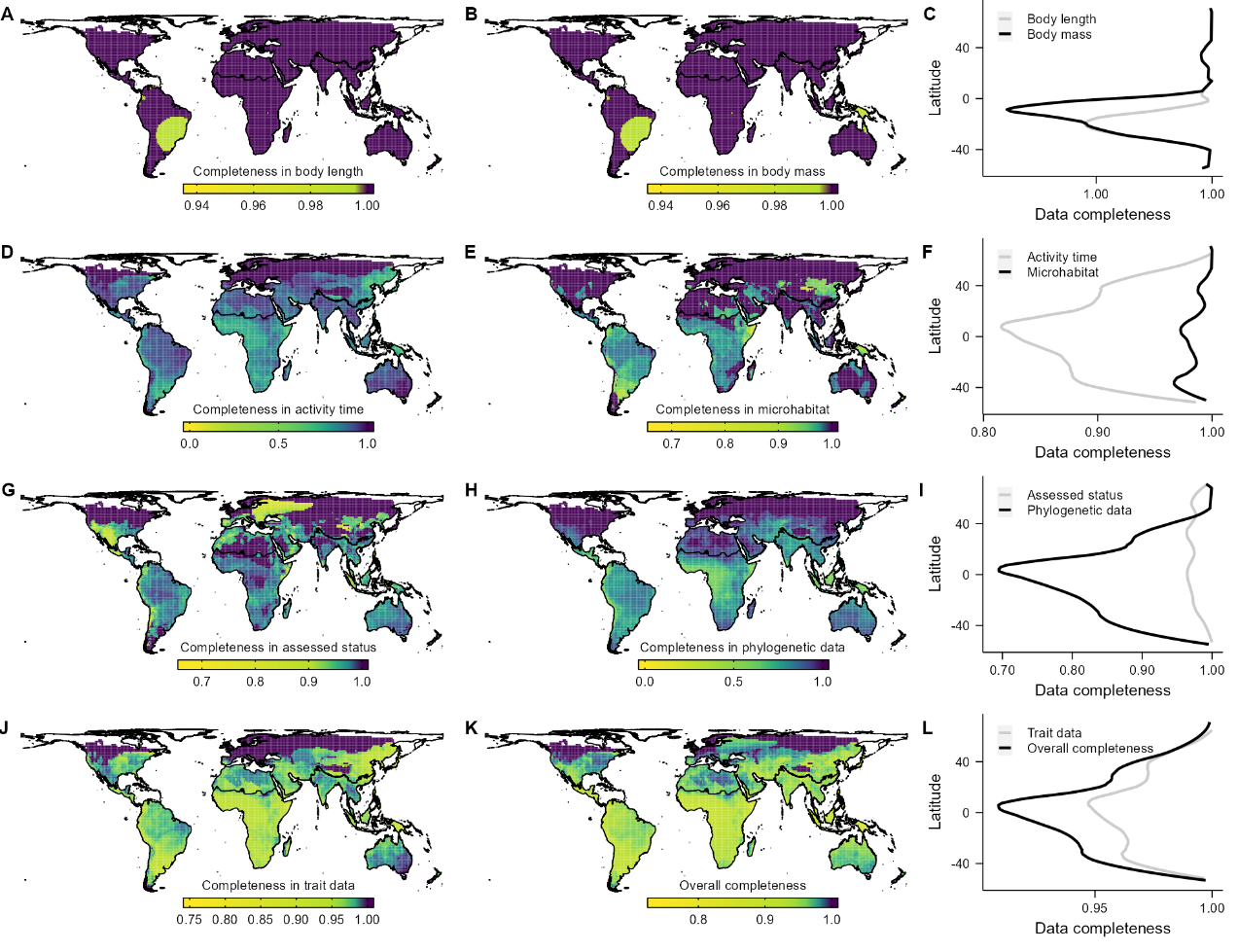

**Figure S6. Average data completeness across squamate assemblages.** Proportion of species with observed values for: (A) body length, (B) body mass, (D) activity time, (E) microhabitat, (G) assessed threat status, (H) phylogenetic data, (J) attribute set, i.e., the average pattern for maps depicted in ABDE, and (K) the complete database, i.e., average pattern for maps depicted in ABDEGH. Maps show grid cell assemblages of 110 × 110 km size in an equal-area projection. Latitudinal plots (C, F, I, L) show the average values of these cells across latitudes. Colour ramps followed Jenks’ natural breaks classification.

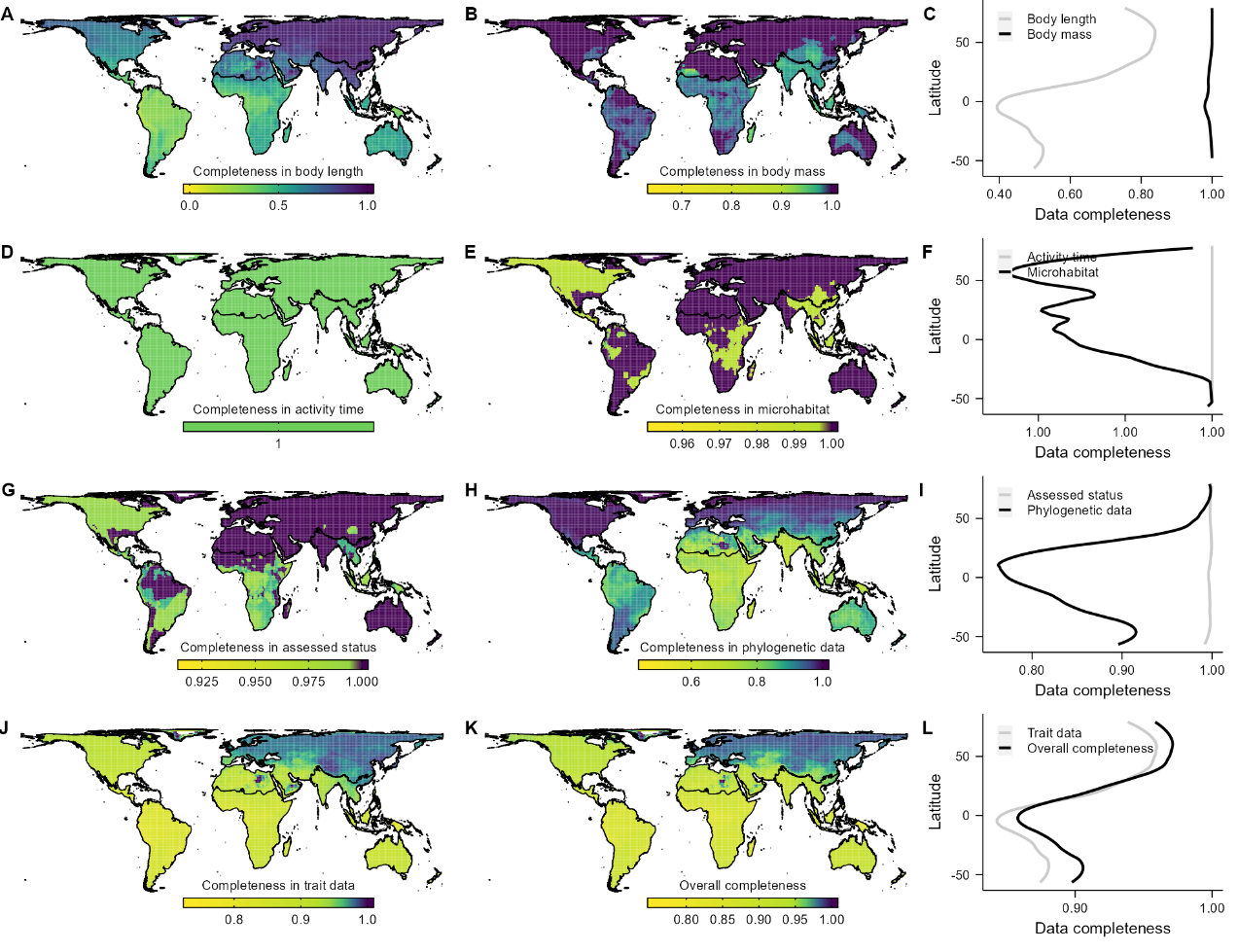

**Figure S7. Average data completeness across bird assemblages.** Proportion of species with observed values for: (A) body length, (B) body mass, (D) activity time, (E) microhabitat, (G) assessed threat status, (H) phylogenetic data, (J) attribute set, i.e., the average pattern for maps depicted in ABDE, and (K) the complete database, i.e., average pattern for maps depicted in ABDEGH. Maps show grid cell assemblages of 110 × 110 km size in an equal-area projection. Latitudinal plots (C, F, I, L) show the average values of these cells across latitudes. Colour ramps followed Jenks’ natural breaks classification.

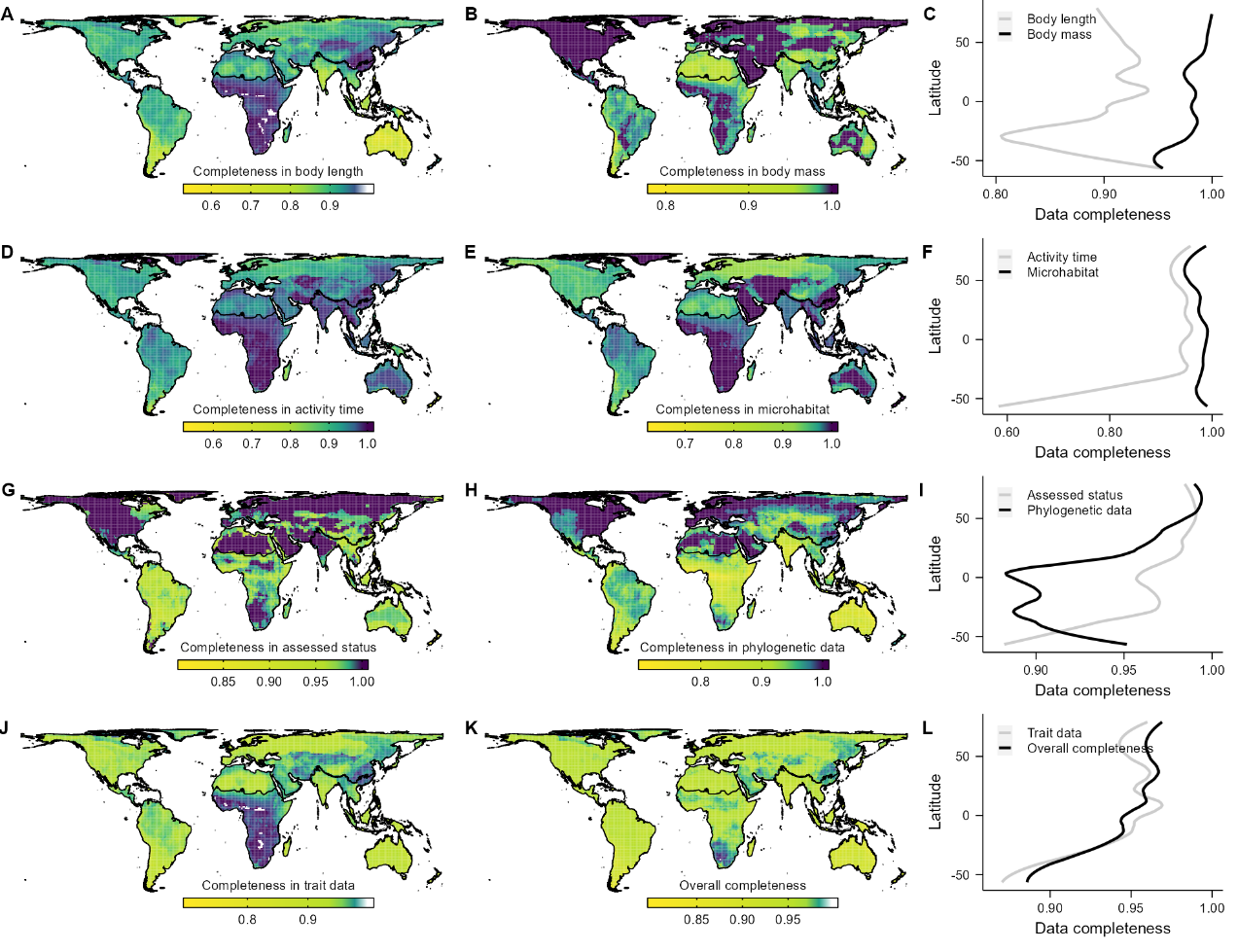

**Figure S8. Average data completeness across mammal assemblages.** Proportion of species with observed values for: (A) body length, (B) body mass, (D) activity time, (E) microhabitat, (G) assessed threat status, (H) phylogenetic data, (J) attribute set, i.e., the average pattern for maps depicted in ABDE, and (K) the complete database, i.e., average pattern for maps depicted in ABDEGH. Maps show grid cell assemblages of 110 × 110 km size in an equal-area projection. Latitudinal plots (C, F, I, L) show the average values of these cells across latitudes. Colour ramps followed Jenks’ natural breaks classification.

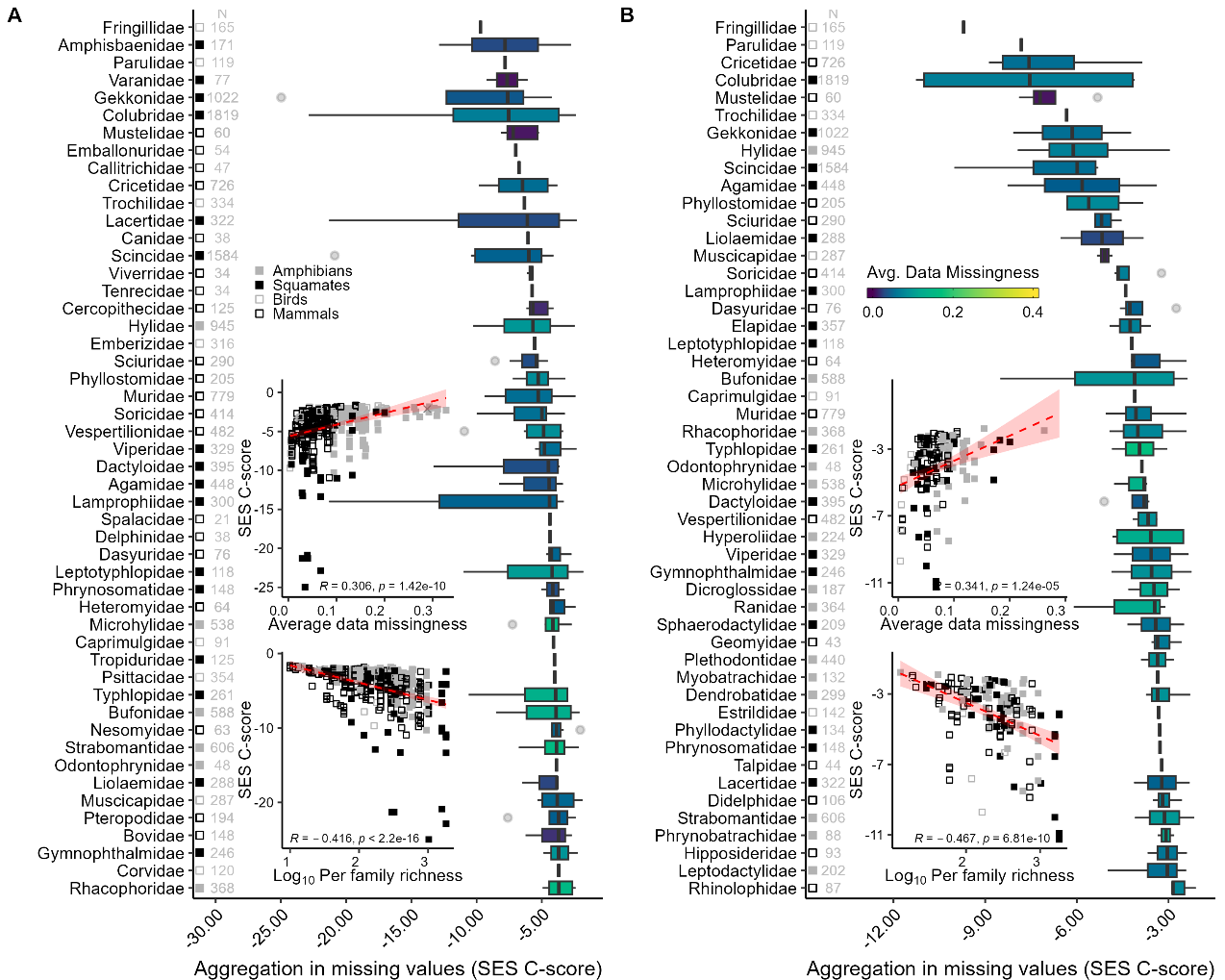

**Figure S9.** **The top 50 tetrapod families with most pronounced patterns of shared data missingness.** For each family, the aggregation metric equals the median value of the standardised effect-size (SES) of the C-score metric computed across (A) all pairwise attribute combinations; or (B) attribute pairs mandatorily involving threat status. Grey numbers on the left side of each panel indicate the per-family species richness.

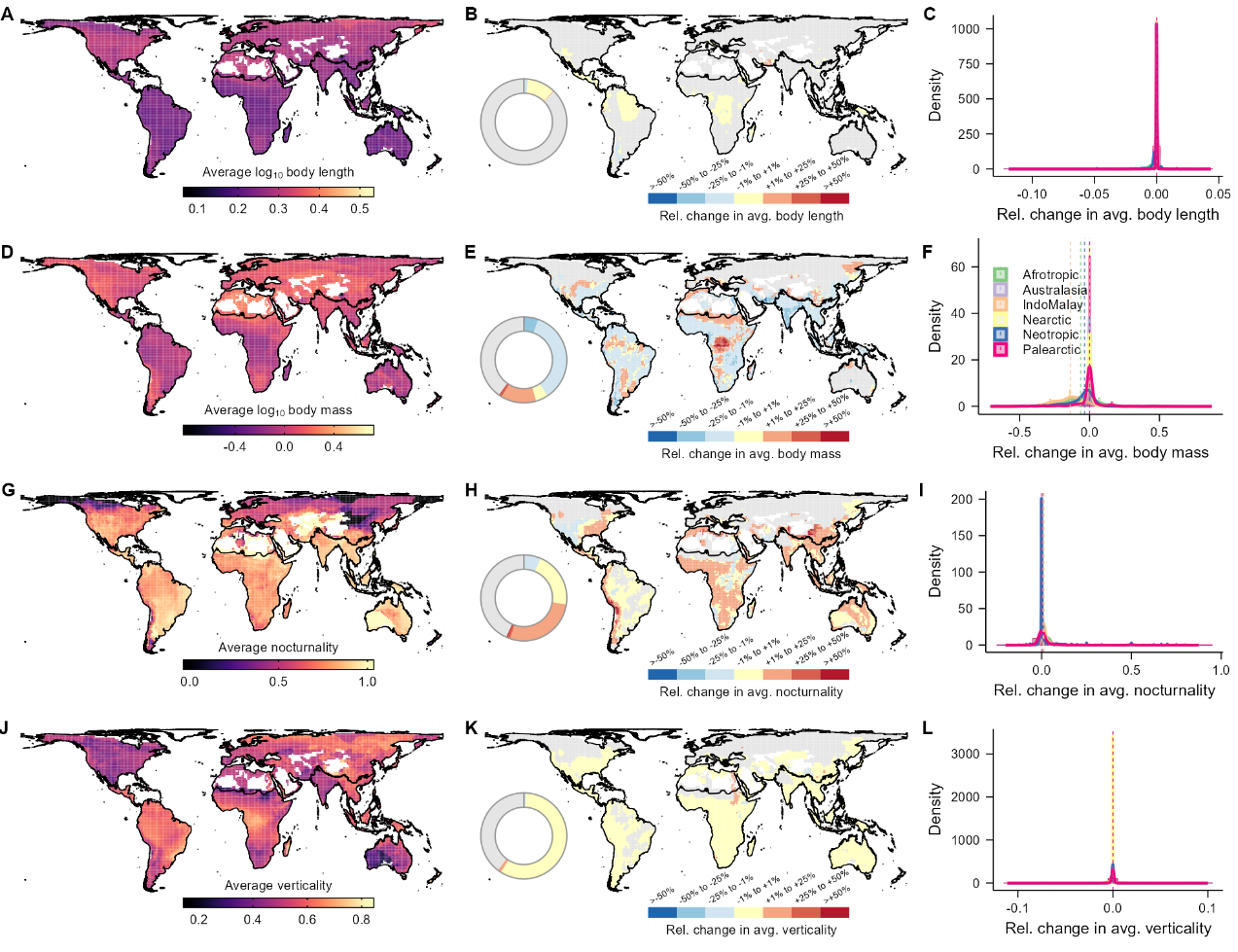

**Figure S10. Changes in average attributes per amphibian assemblage after imputation-based gap-filling.** For each grid cell, maps show the average species attribute value and respective relative change in attribute value after the inclusion of imputed values for (A─C) body length, (D─F) body mass, (G─I) nocturnality, (J─L) verticality. Body length and body mass were log10 transformed before computations. Grey cells indicate assemblages without species with imputed values. Maps draw at the spatial resolution of 110 × 110 km in an equal area projection.

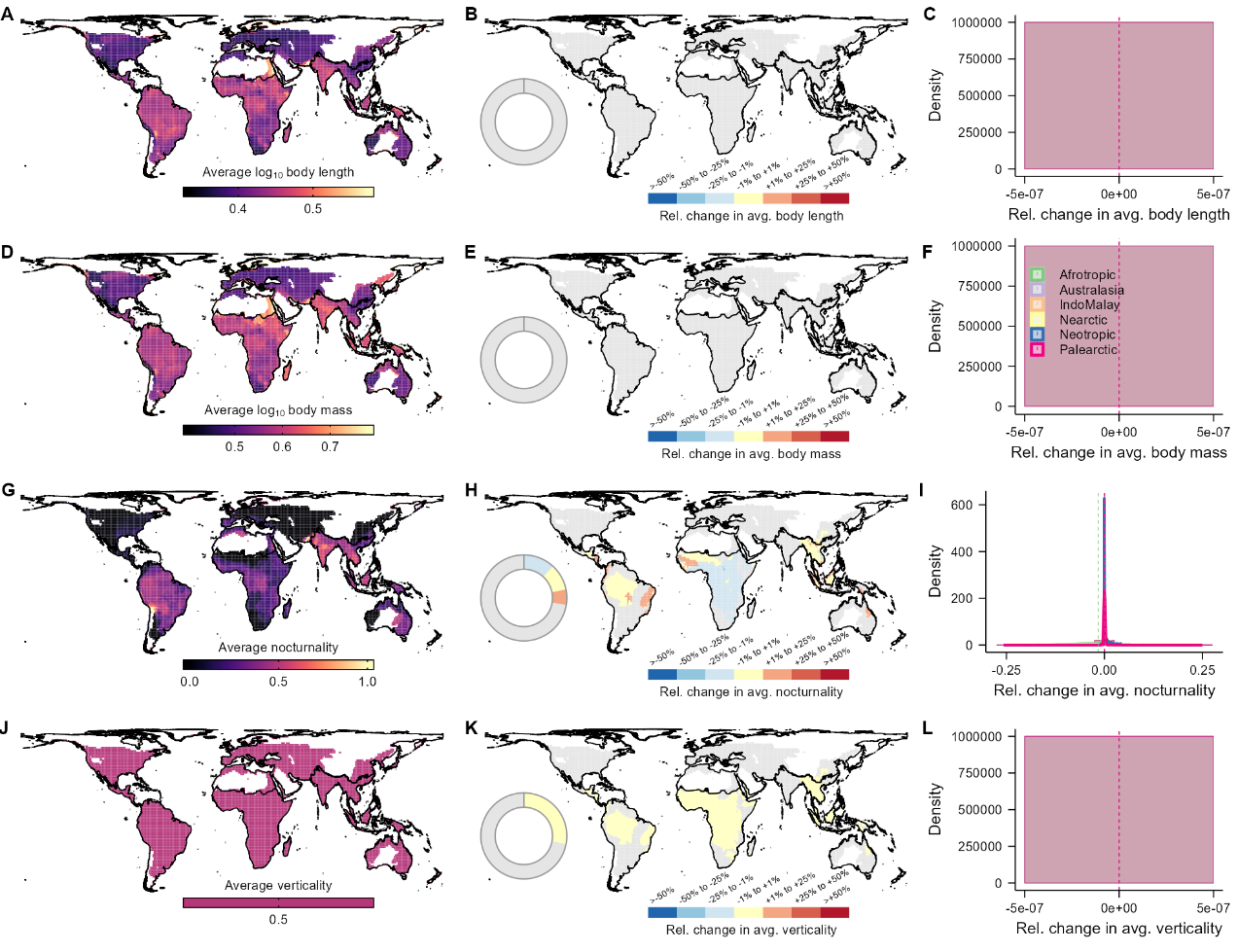

**Figure S11. Changes in average attributes per chelonian and crocodilian assemblage after imputation-based gap-filling.** For each grid cell, maps show the average species attribute value and respective relative change in attribute value after the inclusion of imputed values for (A─C) body length, (D─F) body mass, (G─I) nocturnality, (J─L) verticality. Body length and body mass were log10 transformed before computations. Grey cells indicate assemblages without species with imputed values. Maps draw at the spatial resolution of 110 × 110 km in an equal area projection.

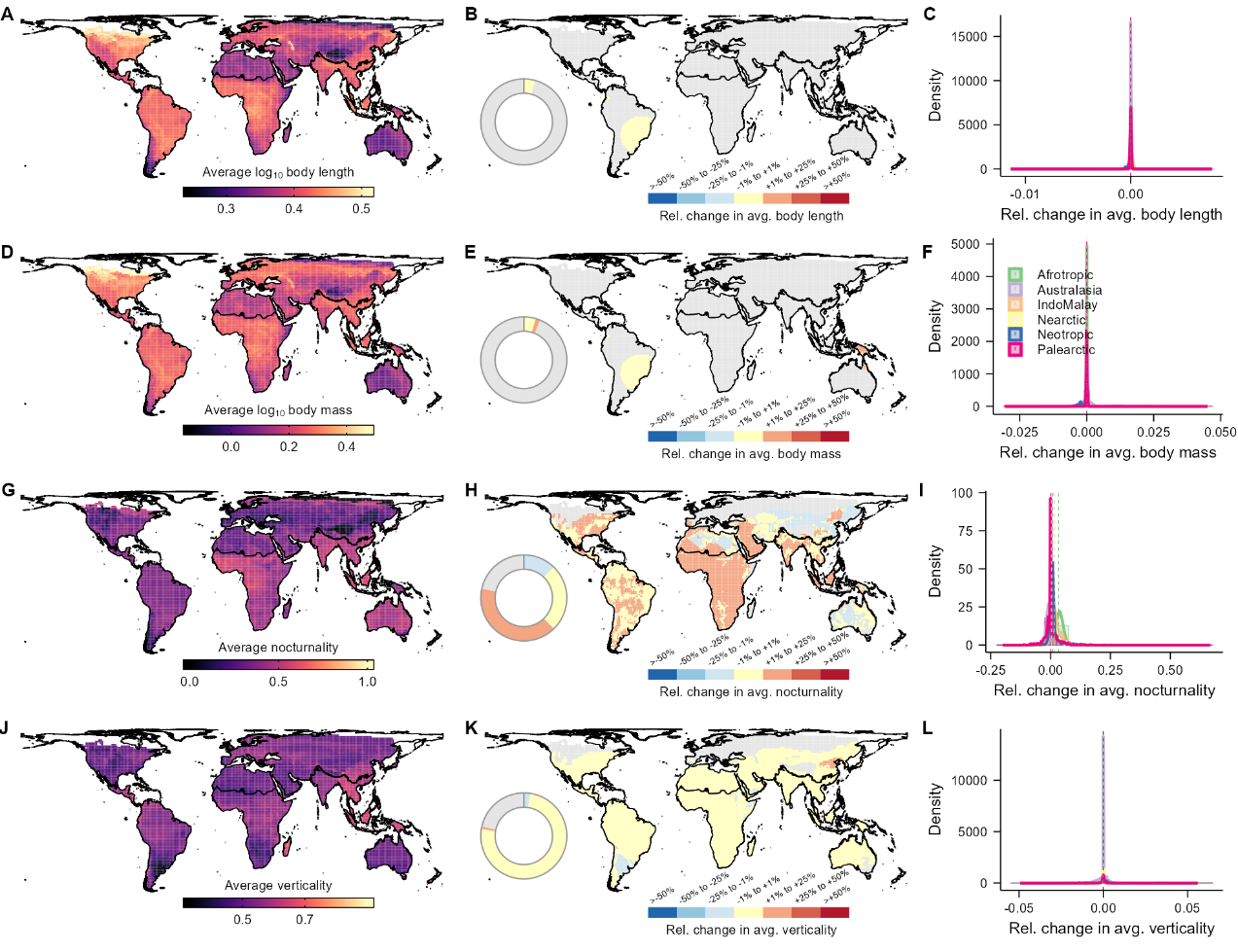

**Figure S12. Changes in average attributes per squamate assemblage after imputation-based gap-filling.** For each grid cell, maps show the average species attribute value and respective relative change in attribute value after the inclusion of imputed values for (A─C) body length, (D─F) body mass, (G─I) nocturnality, (J─L) verticality. Body length and body mass were log10 transformed before computations. Grey cells indicate assemblages without species with imputed values. Maps draw at the spatial resolution of 110 × 110 km in an equal area projection.

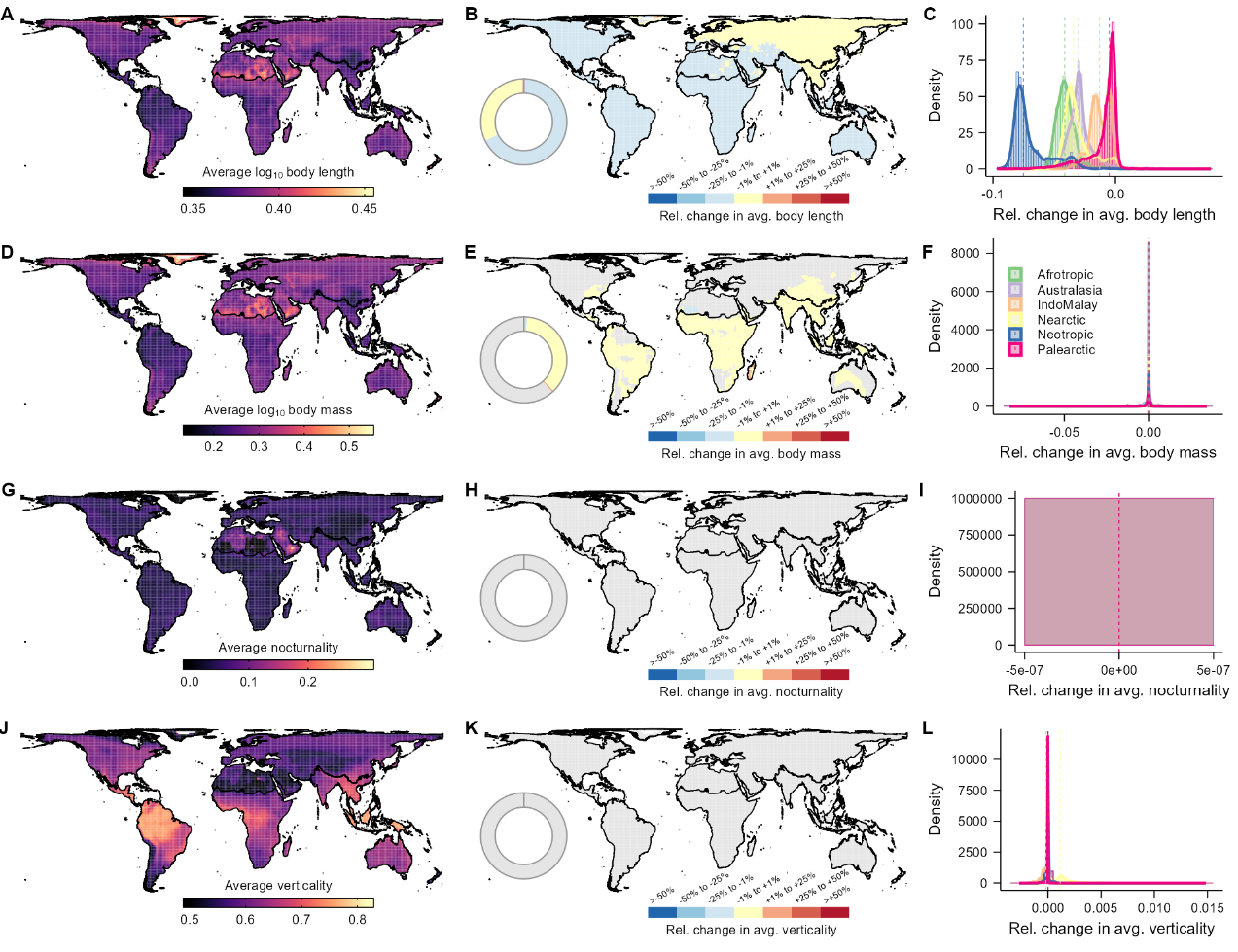

**Figure S13. Changes in average attributes per bird assemblage after imputation-based gap-filling.** For each grid cell, maps show the average species attribute value and respective relative change in attribute value after the inclusion of imputed values for (A─C) body length, (D─F) body mass, (G─I) nocturnality, (J─L) verticality. Body length and body mass were log10 transformed before computations. Grey cells indicate assemblages without species with imputed values. Maps draw at the spatial resolution of 110 × 110 km in an equal area projection.

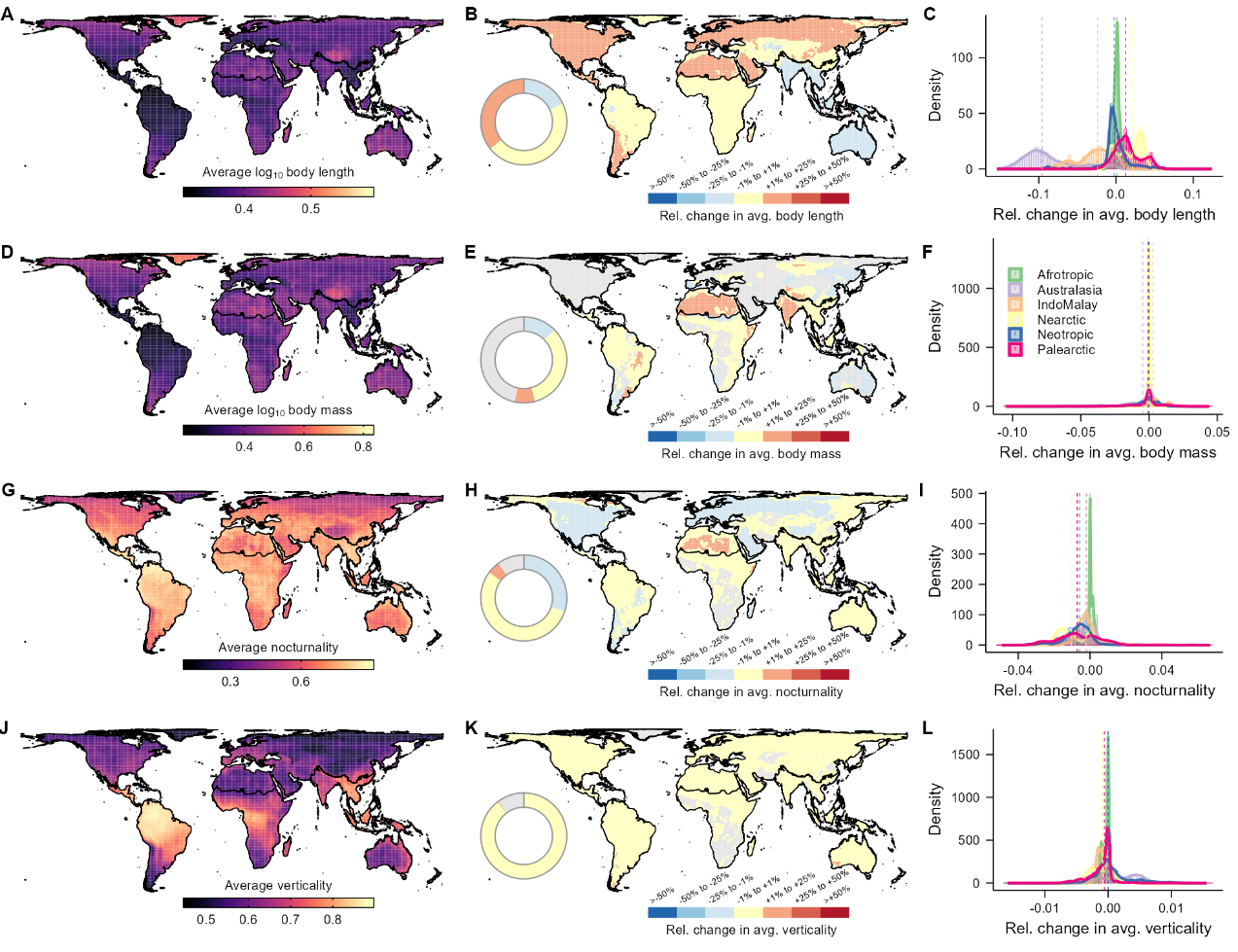

**Figure S14. Changes in average attributes per mammal assemblage after imputation-based gap-filling.** For each grid cell, maps show the average species attribute value and respective relative change in attribute value after the inclusion of imputed values for (A─C) body length, (D─F) body mass, (G─I) nocturnality, (J─L) verticality. Body length and body mass were log10 transformed before computations. Grey cells indicate assemblages without species with imputed values. Maps draw at the spatial resolution of 110 × 110 km in an equal area projection.

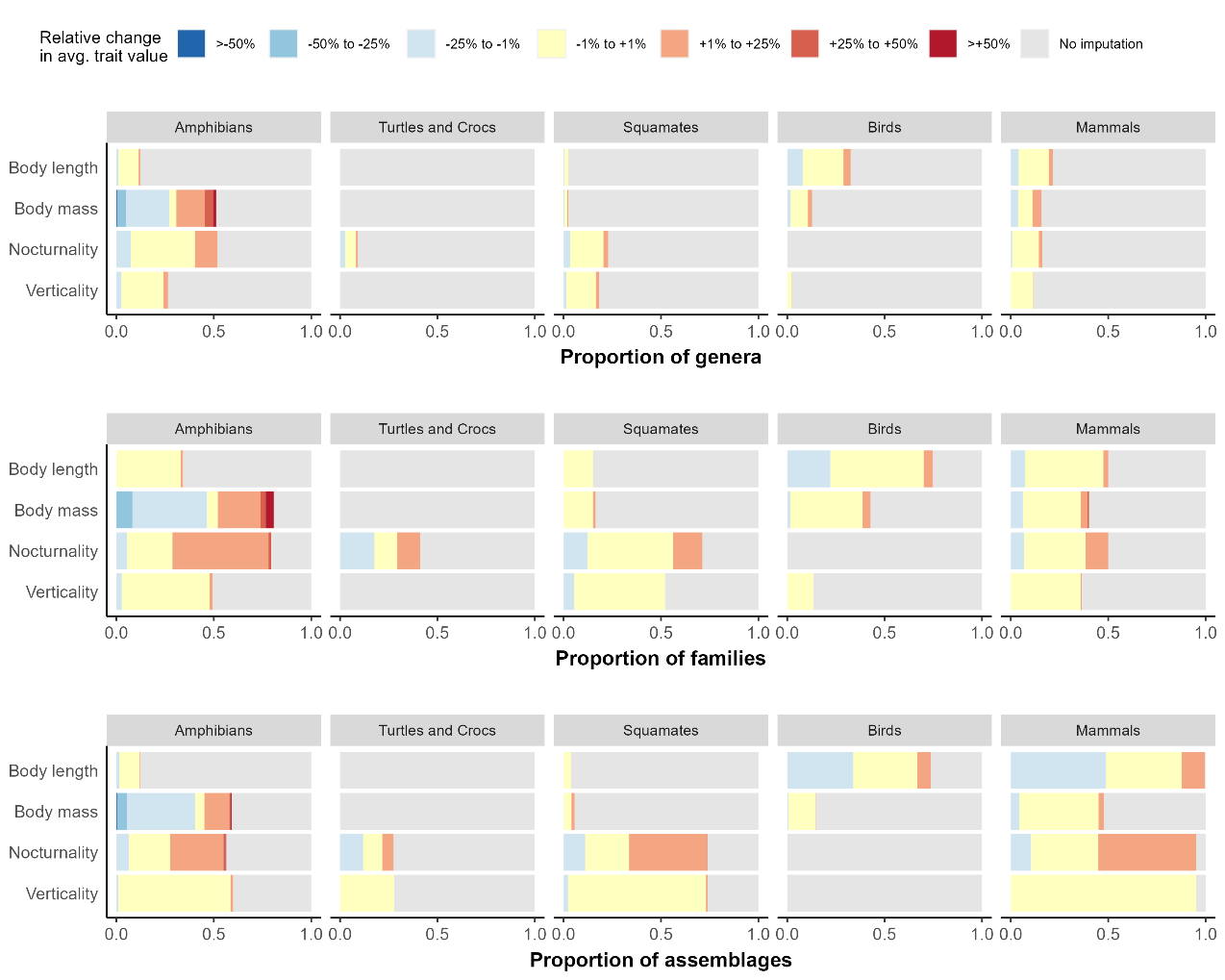

**Figure S15. Proportion of samples across levels of changes in average attributes after imputation-based gap-filling.** Each bar shows the relative number of genera, families, and geographical assemblages facing changes in their average species attribute value after filling missing attributes with imputed values.

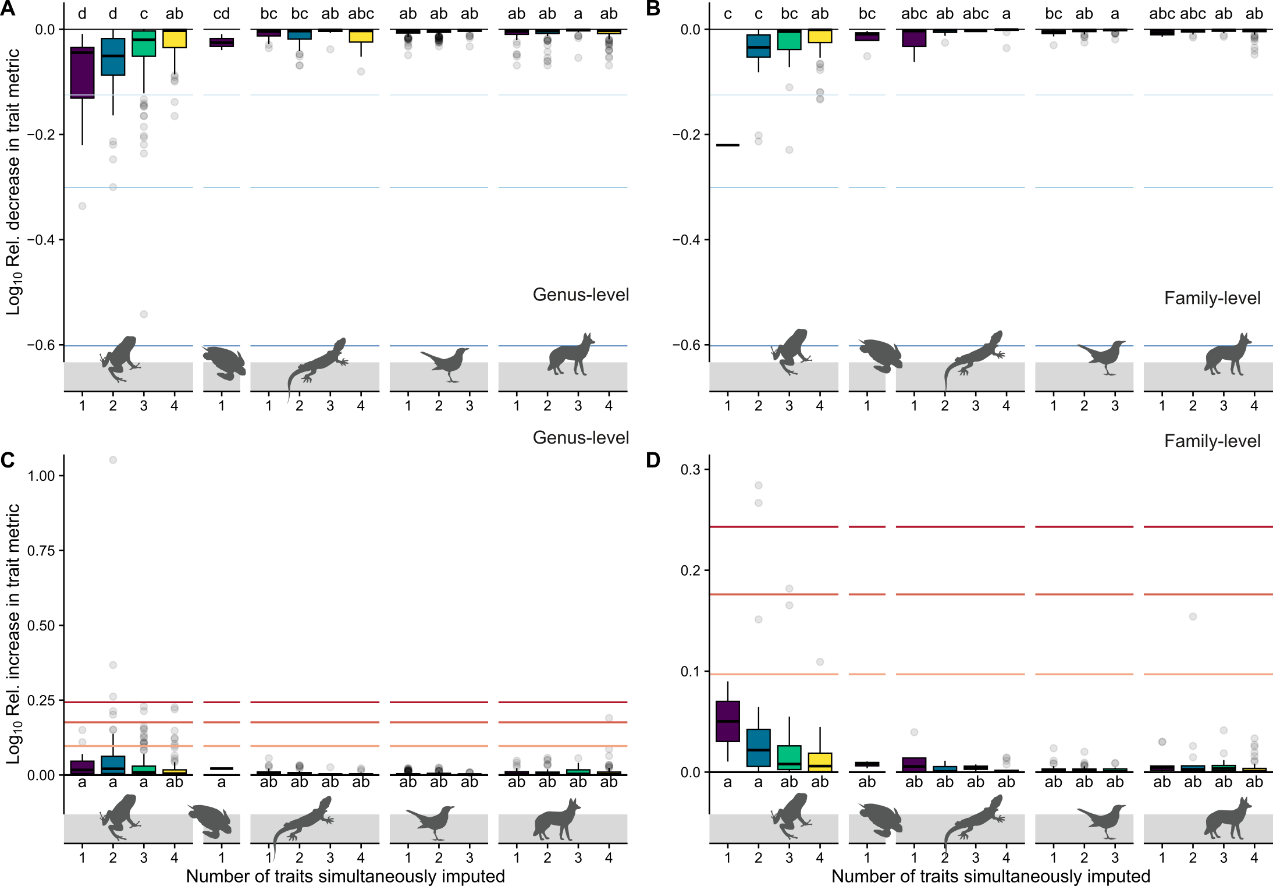

**Figure S16. Relative change in attribute average across increasing number of imputed attributes.** Relative decrease (A, B) or increase (C, D) in attribute metric showed at the (A, C) genus- and (B, D) family-level. Each box denotes the median (horizontal line) and the 25th and 75th percentiles. Vertical lines represent the 95% confidence intervals, and black dots are outliers. Horizontal lines denote the position of 25, 50, 75% of relative decrease (light to dark blue) or increase (light to dark red) in attribute metric. Small capital letters denote the results of the Kruskal–Wallis tests for the difference in medians across relative change in average attribute value.

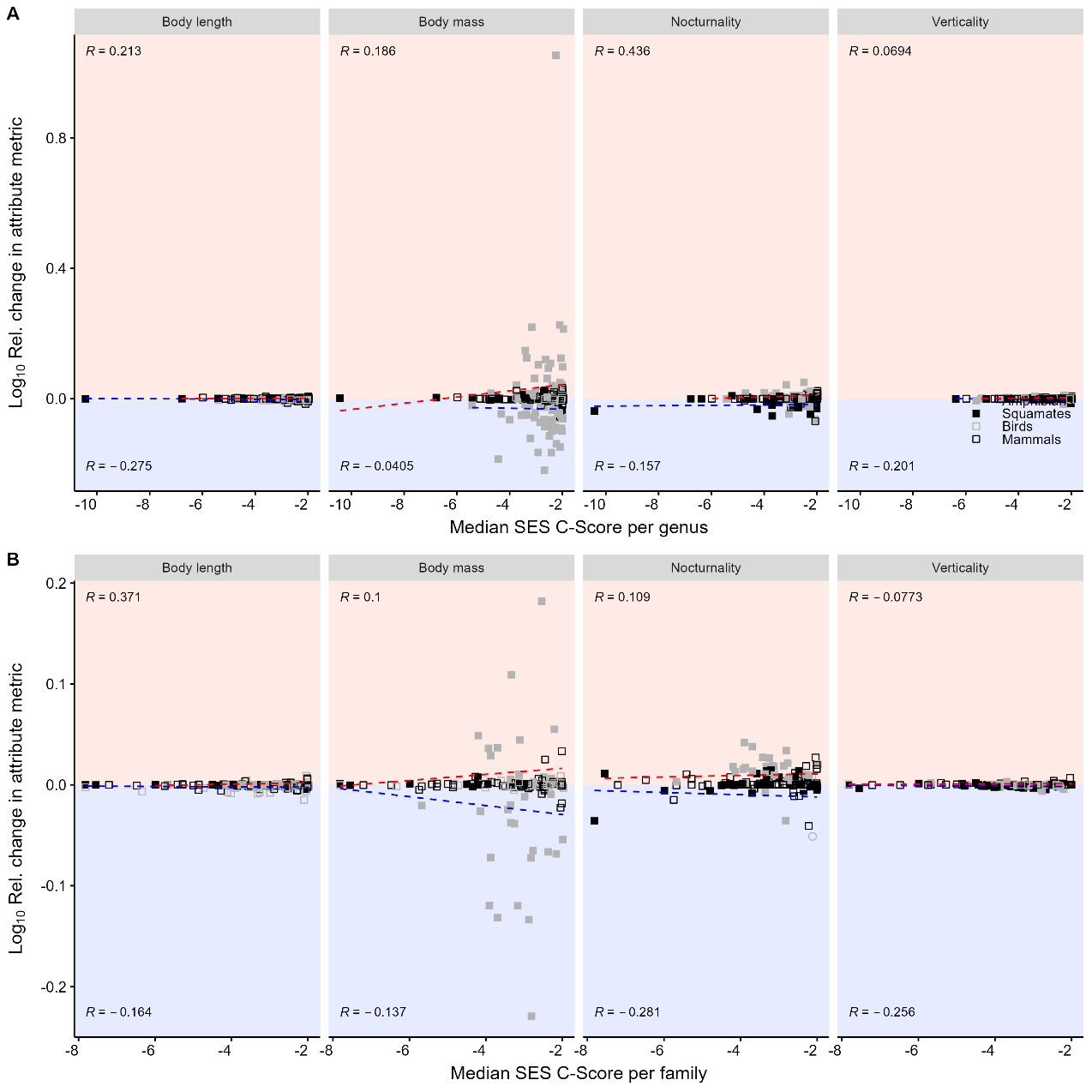

**Figure S17. Changes in aggregate attributes after imputation-based gap-filling in relation to shared missing data.** Relative changes in average attribute value per taxa (geometric mean for body length and mass, and mean for nocturnality and verticality). Each point concerns a taxonomic (A) genus or (B) family. *R* denotes the Spearman correlation coefficient between the relative decrease (blue) or increase (red) in the average attribute value. Only taxa with aggregated patterns of missing data were used in these plots (median Standardised Effect Size of C-score ≤ −1.96). More negative values of SES C-Score indicate a higher degree of shared missing data.

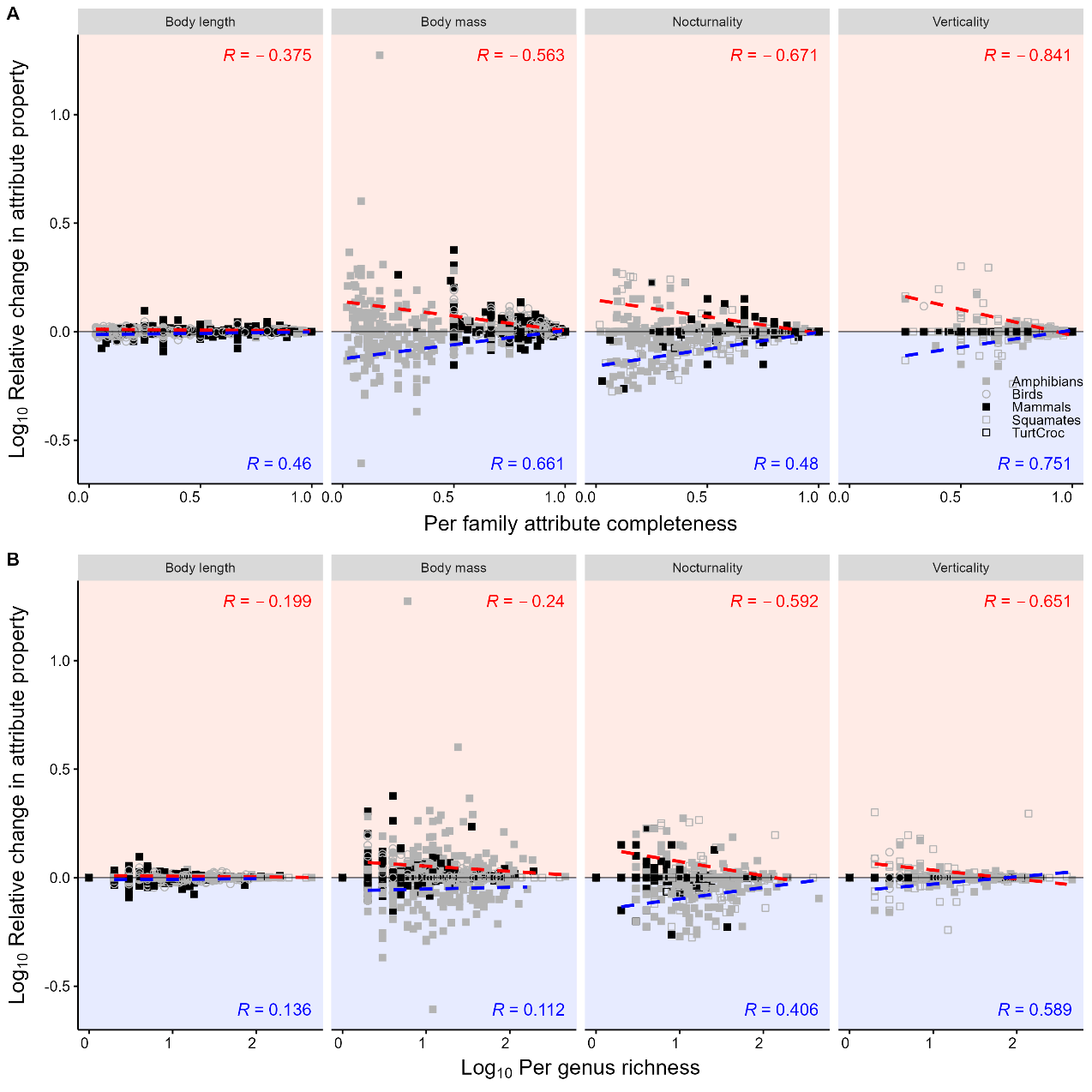

**Figure S18. Changes in aggregate attributes after imputation-based gap-filling in relation to genus completeness and richness.** Relative changes in average attribute value per genus (geometric mean for body length and mass, and mean for nocturnality and verticality). Each point concerns a combination between the relative change in average attribute for a taxonomic genus. *R* denotes the Spearman correlation coefficient between the relative decrease (blue) or increase (red) in the average attribute and (A) per genus attribute completeness and (B) genus richness.

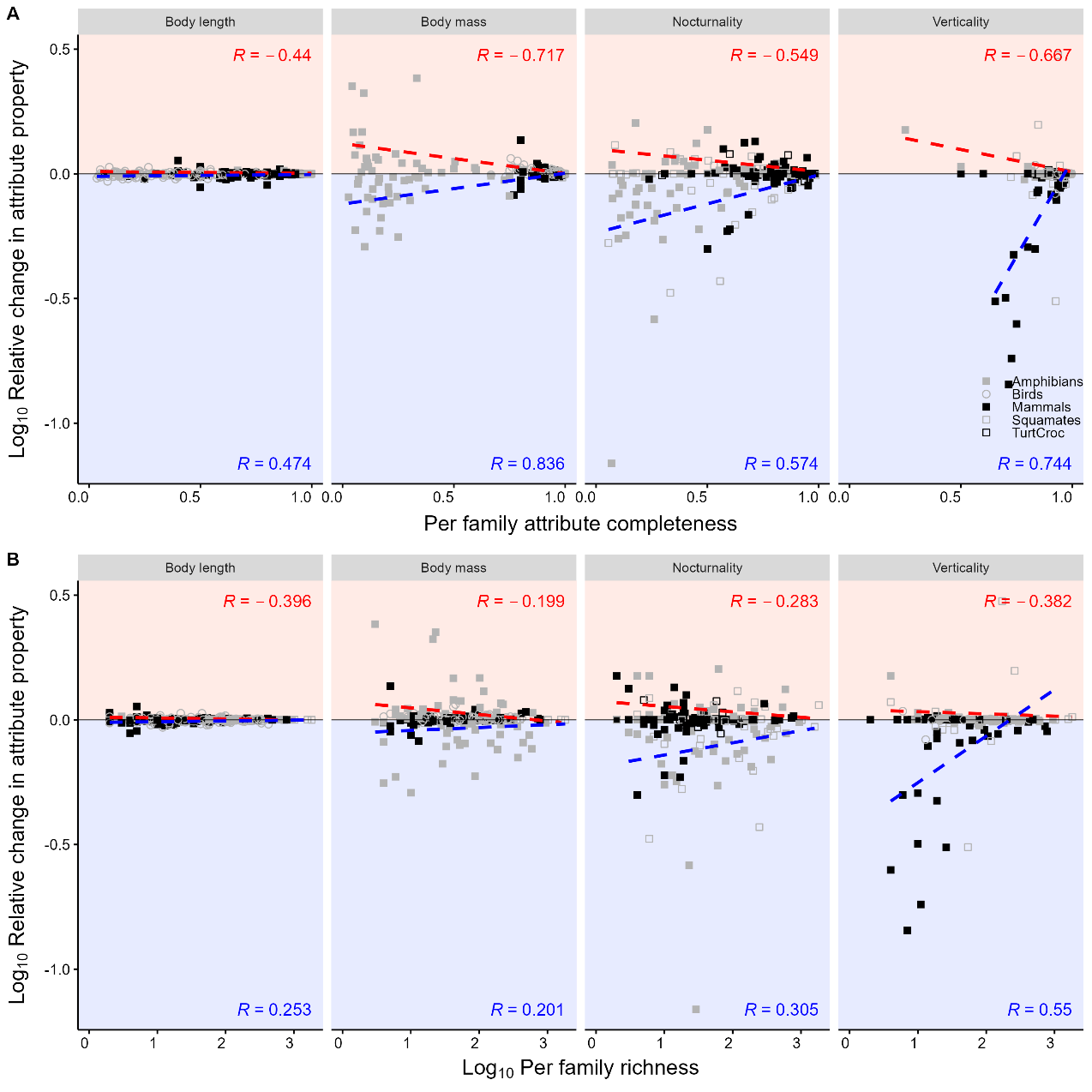

**Figure S19. Changes in aggregate attributes after imputation-based gap-filling in relation to family completeness and richness.** Relative changes in average attribute value per family (geometric mean for body length and mass, and mean for nocturnality and verticality). Each point concerns a combination between the relative change in average attribute for a taxonomic family. *R* denotes the Spearman correlation coefficient between the relative decrease (blue) or increase (red) in the average attribute and (A) per family attribute completeness and (B) family richness.

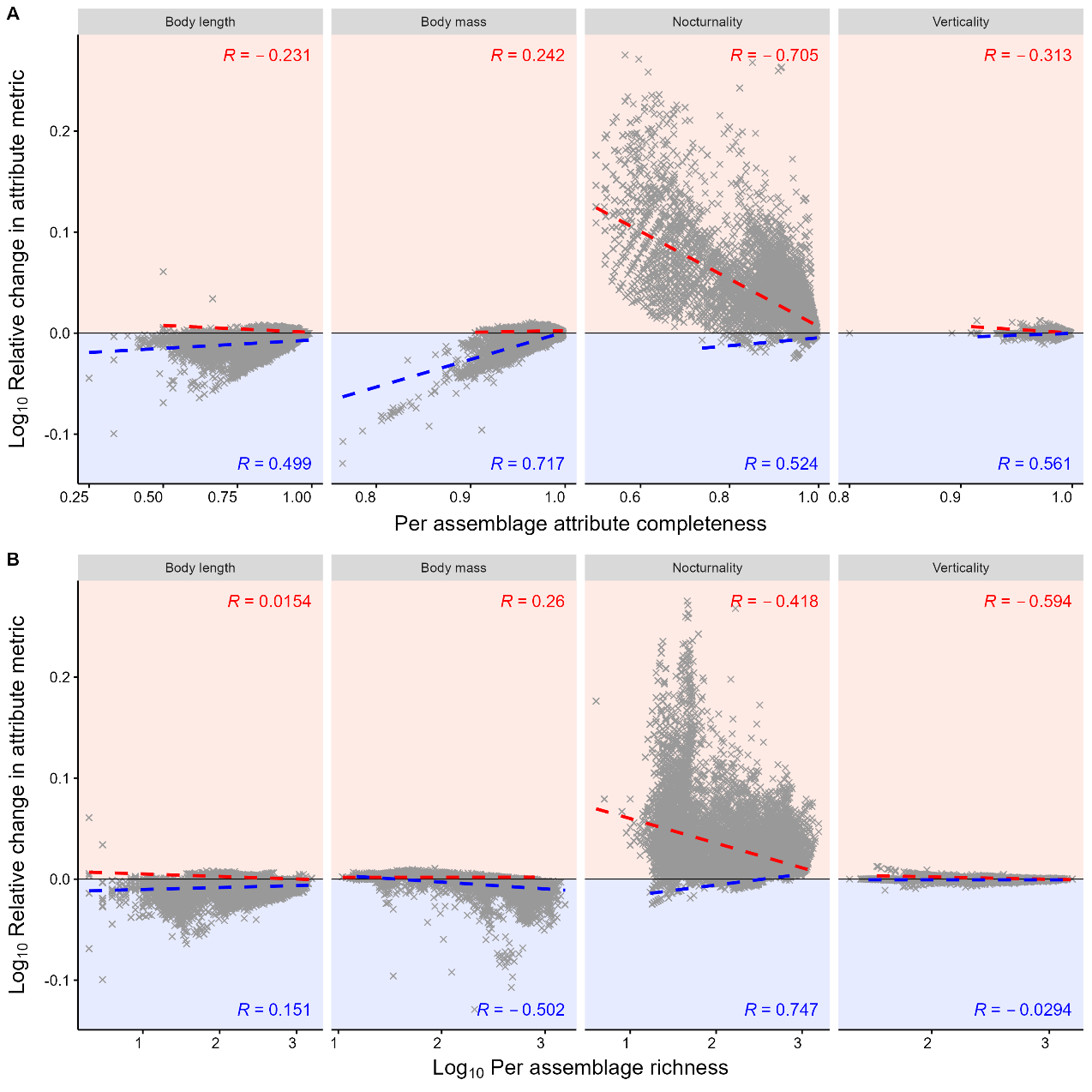

**Figure S20. Changes in aggregate attributes after imputation-based gap-filling in relation to assemblage completeness and richness.** Relative changes in average attribute value per tetrapod assemblage (geometric mean for body length and mass, and mean for nocturnality and verticality. Each point concerns a combination between the relative change in average attribute for a tetrapod assemblage. *R* denotes the Spearman correlation coefficient between the relative decrease (blue) or increase (red) in the average attribute and (A) per assemblage attribute completeness and (B) assemblage richness.

**Supplementary Tables**

Additional tables.

**Table S1.** Variables included in the Tetrapod Traits database.

| **Category** | **Variable name** | **Description** |
| --- | --- | --- |
| Taxonomy | Scientific.Name | Binomial name in the Tetrapod tree. Categorical, 33281 levels. |
|  | Genus | Taxonomic genus. Categorical, 5099 levels. |
|  | Family | Taxonomic family. Categorical, 519 levels. |
|  | Suborder | Taxonomic suborder or similar higher-level taxonomic rank. Categorical, 88 levels. |
|  | Order | Taxonomic order. Categorical, 74 levels. |
|  | Class | Taxonomic class. Categorical, 4 levels. |
|  | Authority | Authority name associated with the scientific name. |
|  | YearOfDescription | Year in which the species was first described. |
| Tree | TreeTaxon | Tree-taxonomy followed. Categorical, 5 levels. |
|  | TreeImputed | Phylogenetic relationship imputed in the fully-sampled trees. Binary: 1 = Imputed, 0 = Not imputed. |
| Body Size | BodyLength_mm | Maximum body length, in millimetres (mm). |
|  | LengthMeasure | Body length measure. Categorical, 4 levels: CL = carapace length, SVL = snout-vent length, TL = total length, HBL = head-body length. |
|  | ImputedLength | Body length value imputed. Binary: 1 = Imputed, 0 = Not imputed. |
|  | SourceBodyLength | Source for body length data. |
|  | BodyMass_g | Maximum body mass in grams (g). |
|  | ImputedMass | Body mass value imputed. Binary: 1 = Imputed, 0 = Not imputed. |
|  | SourceBodyMass | Source for body mass data. |
| Activity time | Diu | Diurnal activity. Binary: 1 = Present, 0 = Absent. |
|  | Noc | Nocturnal activity. Binary: 1 = Present, 0 = Absent. |
|  | ImputedActTime | Activity Time value imputed. Binary: 1 = Imputed, 0 = Not imputed. |
|  | SourceActTime | Source for activity time data. |
|  | Nocturnality | Nocturnality score. Numeric: 0 = Diurnal, 0.5 = Cathemeral/Crepuscular, 1 = Nocturnal. |
| Microhabitat | Fos | Fossorial microhabitat use. Binary: 1 = Present, 0 = Absent. |
|  | Ter | Terrestrial microhabitat use. Binary: 1 = Present, 0 = Absent. |
|  | Aqu | Aquatic microhabitat use. Binary: 1 = Present, 0 = Absent. |
|  | Arb | Arboreal microhabitat use. Binary: 1 = Present, 0 = Absent. |
|  | Aer | Aerial microhabitat use. Binary: 1 = Present, 0 = Absent. |
|  | ImputedHabitat | Microhabitat value imputed. Binary: 1 = Imputed, 0 = Not imputed. |
|  | SourceHabitat | Source for microhabitat data. |
|  | Verticality | Verticality score. Numeric: 0 = fossorial, 0.25 = semifossorial, 0.5 = terrestrial or aquatic, 0.75 = semiarboreal, 1 = arboreal or aerial. |
| Macrohabitat | MajorHabitat_1 | Species major habitat includes Forest. Binary: 1 = Yes, 0 = No. |
|  | MajorHabitat_2 | Species major habitat includes Savanna. Binary: 1 = Yes, 0 = No. |
|  | MajorHabitat_3 | Species major habitat includes Shrubland. Binary: 1 = Yes, 0 = No. |
|  | MajorHabitat_4 | Species major habitat includes Grassland. Binary: 1 = Yes, 0 = No. |
|  | MajorHabitat_5 | Species major habitat includes Wetlands (inland). Binary: 1 = Yes, 0 = No. |
|  | MajorHabitat_6 | Species major habitat includes Rocky Areas. Binary: 1 = Yes, 0 = No. |
|  | MajorHabitat_7 | Species major habitat includes Caves & Subterranean Habitats. Binary: 1 = Yes, 0 = No. |
|  | MajorHabitat_8 | Species major habitat includes Desert. Binary: 1 = Yes, 0 = No. |
|  | MajorHabitat_9 | Species major habitat includes Marine Neritic. Binary: 1 = Yes, 0 = No. |
|  | MajorHabitat_10 | Species major habitat includes Marine Oceanic. Binary: 1 = Yes, 0 = No. |
|  | MajorHabitat_12 | Species major habitat includes Marine Intertidal. Binary: 1 = Yes, 0 = No. |
|  | MajorHabitat_13 | Species major habitat includes Marine Coastal/Supratidal. Binary: 1 = Yes, 0 = No. |
|  | MajorHabitat_14 | Species major habitat includes Artificial Terrestrial. Binary: 1 = Yes, 0 = No. |
|  | MajorHabitat_15 | Species major habitat includes Artificial Aquatic. Binary: 1 = Yes, 0 = No. |
|  | MajorHabitat_16 | Species major habitat includes Introduced Vegetation. Binary: 1 = Yes, 0 = No. |
|  | MajorHabitat_17 | Species major habitat includes Other. Binary: 1 = Yes, 0 = No. |
|  | MajorHabitatSum | Number of major habitats in which the species occur. |
|  | ImputedMajorHabitat | Major Habitat value taxonomically imputed. Binary: 1 = Imputed, 0 = Not imputed. |
|  | SourceMajorHabitat | Source for major habitat data. |
| Ecosystem | EcoTer | Species occurs in terrestrial ecosystem. Binary: 1 = Yes, 0 = No. |
|  | EcoFresh | Species occurs in freshwater ecosystem. Binary: 1 = Yes, 0 = No. |
|  | EcoMar | Species occurs in marine ecosystem. Binary: 1 = Yes, 0 = No. |
|  | EcoSystemSum | Number of ecosystems in which the species occur. |
|  | ImputedEcosystem | Ecosystem value taxonomically imputed. Binary: 1 = Imputed, 0 = Not imputed. |
|  | SourceEcosystem | Source for ecosystem data. |
| Threat Status | IUCN_Binomial | Binomial name spelling according to the IUCN red list v. 2022-2 (if available). |
|  | AssessedStatus | Assessed threat statuses from IUCN Red List. |
|  | SourceStatus | IUCN Red List version consulted for the assessed status. |
| Geography | RangeSize | Number of 110×110 km equal-area grid cells occupied by the species geographic range. |
|  | Longitude | Average within-range longitude in decimal degrees. |
|  | Latitude | Average within-range latitude in decimal degrees. |
|  | Afrotropic | Proportion of the species geographic range within the Afrotropical realm. |
|  | Australasia | Proportion of the species geographic range within the Australasia realm. |
|  | IndoMalay | Proportion of the species geographic range within the IndoMalay realm. |
|  | Afrotropic | Proportion of the species geographic range within the Afrotropical realm. |
|  | Australasia | Proportion of the species geographic range within the Australasia realm. |
|  | IndoMalay | Proportion of the species geographic range within the IndoMalay realm. |
|  | Nearctic | Proportion of the species geographic range within the Nearctic realm. |
|  | Neotropic | Proportion of the species geographic range within the Neotropic realm. |
|  | Oceania | Proportion of the species geographic range within the Oceania realm. |
|  | Palearctic | Proportion of the species geographic range within the Palearctic realm. |
|  | Antarctic | Proportion of the species geographic range within the Antarctic realm. |
|  | Insularity | Species is insular endemic. Binary: 1 = Yes, 0 = No. |
|  | SourceInsularity | Source for insularity data. |
| Environment | AnnuMeanTemp | Average within-range annual mean temperature (Celsius degree). Data derived from CHELSA v. 1.2 [1]. |
|  | AnnuPrecip | Average within-range annual precipitation (mm). Data derived from CHELSA v. 1.2 [1]. |
|  | TempSeasonality | Average within-range temperature seasonality (Standard deviation × 100). Data derived from CHELSA v. 1.2 [1]. |
|  | PrecipSeasonality | Average within-range precipitation seasonality (Coefficient of Variation). Data derived from CHELSA v. 1.2 [1]. |
|  | Elevation | Average within-range elevation (metres). Data derived from topographic layers in EarthEnv [2]. |
| Human Influence | ETA50K | Average within-range estimated time to travel to cities with a population >50K in the year 2015. Data from [3]. |
|  | HumanDensity | Average within-range human population density in year 2017. Data derived from HYDE v. 3.2 [4]. |
|  | PropUrbanArea | Proportion of species range map covered by built-up area, such as towns, cities, etc. at year 2017 [4]. |
|  | PropCroplandArea | Proportion of species range map covered by cropland area, identical to FAO's category ‘Arable land and permanent crops’ at year 2017 [4]. |
|  | PropPastureArea | Proportion of species range map covered by cropland, defined as Grazing land with an aridity index > 0.5, assumed to be more intensively managed (converted in climate models) at year 2017 [4]. |
|  | PropRangelandArea | Proportion of species range map covered by rangeland, defined as Grazing land with an aridity index < 0.5, assumed to be less or not managed (not converted in climate models) at year 2017 [4]. |

1. Karger DN, Conrad O, Böhner J, Kawohl T, Kreft H, Soria-Auza RW, et al. Climatologies at high resolution for the earth’s land surface areas. Sci Data. 2017;4: 170122. doi:10.1038/sdata.2017.122

2. Amatulli G, Domisch S, Tuanmu M-N, Parmentier B, Ranipeta A, Malczyk J, et al. A suite of global, cross-scale topographic variables for environmental and biodiversity modeling. Sci Data. 2018;5: 180040. doi:10.1038/sdata.2018.40

3. Nelson A. Estimated travel time to the nearest city of 50,000 or more people in year 2000. Ispra: Global Environment Monitoring Unit - Joint Research Centre of the European Commission; 2008. Available: http://forobs.jrc.ec.europa.eu/products/gam/

4. Klein Goldewijk K, Beusen A, Doelman J, Stehfest E. Anthropogenic land use estimates for the Holocene-HYDE 3.2. Earth Syst Sci Data. 2017;9: 927–953. doi:10.5194/essd-9-927-2017

**Table S2.** Number of species with natural history data available per tetrapod group.

| **Attribute/Group** | **Obs. data** | **Taxonomic Imputation** | **Phylogenetic Imputation** | **Completeness Pre-Imputing** |
| --- | --- | --- | --- | --- |
| **Amphibians (7,238 spp.)** |  |  |  |  |
| Body length^*^ | 7083 | - | 155 | 97.9% |
| Body mass^*^ | 1547 | - | 5691 | 21.4% |
| Activity time^*^ | 3501 | 183 | 3554 | 48.4% |
| Microhabitat^*^ | 6669 | 453 | 116 | 92.1% |
| Macrohabitat | 6945 | 291 | - | 95.9% |
| Ecosystem | 7092 | 146 | - | 98.0% |
| Insularity | 7238 | - | - | 100% |
| Non-DD threat status | 6388 | - | - | 88.3% |
| Expert-based range map | 7236 | - | - | >99.9% |
| **Chelonians and crocodilians (384 spp.)** |  |  |  |  |
| Body length^*^ | 384 | - | - | 100% |
| Body mass^*^ | 384 | - | - | 100% |
| Activity time^*^ | 331 | 1 | 52 | 86.2% |
| Microhabitat^*^ | 384 | - | - | 100% |
| Macrohabitat | 344 | 20 | - | 89.6% |
| Ecosystem | 384 | - | - | 100% |
| Insularity | 384 | - | - | 100% |
| Non-DD threat status | 290 | - | - | 75.5% |
| Expert-based range map | 384 | - | - | 100% |
| **Squamates (9,755 spp.)** |  |  |  |  |
| Body length^*^ | 9728 | - | 27 | 99.7% |
| Body mass^*^ | 9727 | - | 28 | 99.7% |
| Activity time^*^ | 6778 | 429 | 2548 | 69.5% |
| Microhabitat^*^ | 9162 | 333 | 260 | 93.9% |
| Macrohabitat | 9323 | 271 | - | 95.6% |
| Ecosystem | 9504 | 250 | - | 97.4% |
| Insularity | 9754 | - | - | >99.9% |
| Non-DD threat status | 8146 | - | - | 83.5% |
| Expert-based range map | 9747 | - | - | 99.9% |
| **Birds (9,993 spp)** |  |  |  |  |
| Body length^*^ | 3161 | - | 6832 | 31.6% |
| Body mass^*^ | 9436 | - | 557 | 94.4% |
| Activity time^*^ | 9993 | - | - | 100% |
| Microhabitat^*^ | 9927 | 66 | - | 99.3% |
| Macrohabitat | 9922 | 71 | - | 99.3% |
| Ecosystem | 9880 | 113 | - | 98.9% |
| Insularity | 9993 | - | - | 100% |
| Non-DD threat status | 9933 | - | - | 99.4% |
| Expert-based range map | 9993 | - | - | 100% |
| **Mammals (5,911 spp.)** |  |  |  |  |
| Body length^*^ | 4802 | - | 1109 | 81.2% |
| Body mass^*^ | 5435 | - | 476 | 91.9% |
| Activity time^*^ | 5056 | 253 | 602 | 85.5% |
| Microhabitat^*^ | 5584 | 258 | 69 | 94.5% |
| Macrohabitat | 5756 | 119 | - | 97.4% |
| Ecosystem | 5808 | 102 | - | 98.3% |
| Insularity | 5911 | - | - | 100% |
| Non-DD threat status | 5045 | - | - | 85.3% |
| Expert-based range map | 5909 | - | - | >99.9% |

^*^ Attribute with 100% of completeness post-imputation.
